# Exploring DNA structures in real-time polymerase kinetics using Pacific Biosciences sequencer data

**DOI:** 10.1101/001024

**Authors:** Sterling Sawaya, James Boocock, Michael A. Black, Neil Gemmell

## Abstract

Pausing of DNA polymerase can indicate the presence of a DNA structure that differs from the canonical double-helix. Here we detail a method to investigate how polymerase pausing in the Pacific Biosciences sequencer reads can be related to DNA structure. The Pacific Biosciences sequencer uses optics to view a polymerase and its interaction with a single DNA molecule in real-time, offering a unique way to detect potential alternative DNA structures. We have developed a new way to examine polymerase kinetics and relate it to the DNA sequence by using a wavelet transform of read information from the sequencer. We use this method to examine how polymerase kinetics are related to nucleotide base composition. We then examine tandem repeat sequences known for their ability to form different DNA structures: (CGG)n and (CG)n repeats which can, respectively, form G-quadruplex DNA and Z-DNA. We find pausing around the (CGG)n repeat that may indicate the presence of G-quadruplexes in some of the sequencer reads. The (CG)n repeat does not appear to cause polymerase pausing, but its kinetics signature nevertheless suggests the possibility that alternative nucleotide conformations may sometimes be present. We discuss the implications of using our method to discover DNA sequences capable of forming alternative structures. The analyses presented here can be reproduced on any Pacific Biosciences kinetics data for any DNA pattern of interest using an R package that we have made publicly available.

**Author Summary:** DNA can be found in various forms that differ from the double-helix first discovered by Watson and Crick in 1953. These alternative DNA structures depend on the DNA sequence, and researchers continue to explore which sequences have the potential to form alternative structures. Here we advance the use of Pacific Biosciences sequencer data to explore potential alternative DNA structures. The Pacific Biosciences sequencer provides an unprecedented way to examine the interaction between DNA polymerase and DNA by following a single polymerase in real time as it copies a DNA molecule. The pausing of DNA polymerase is a common method for exploring the DNA sequences that have the potential to form alternative DNA structures, and Pacific Biosciences data has previously been used to measure polymerase pausing at a slipped strand structure. DNA polymerase is known to pause at some of these alternative structures, such as the structure known as the G-quadruplex, a DNA structure that has potentially importing regulatory significance. We examine polymerase kinetics around a G-quadruplex, and find evidence of polymerase pausing in the Pacific Biosciences kinetics. We provide a method, with publicly available code, so that others can examine these polymerase kinetics for any sequence of interest.

## Introduction

The primary structure of DNA was first discovered in 1953 by Watson and Crick as a right-handed double helix [1], commonly referred to as B-DNA. Since then, other DNA structures have been discovered, some with potential biological significance. These structures include the left-handed double helix, Z-DNA, the triple-helix, H-DNA, slipped-strand hairpin structures, cruciforms, I-motifs and G-quadruplexes (reviewed in [2]). The biological significance of these structures remains disputed, but compelling evidence for the biological function of two of these structures exists: Z-DNA, which is associated with transcription and chromatin remodeling [3–7], and more recently, G-quadruplex DNA which was visualized in living cells and was associated with DNA synthesis [8].

Some of these non-B-DNA structures can be problematic for polymerase-chain-reactions (PCR, [9]). Non-B-DNA structures can interfere with polymerase, resulting in a reduced rate of DNA synthesis [9–15]. Polymerase impedance can introduce genotyping errors because amplification biases can cause some alleles to be lost during amplification, termed “allelic drop-out” [9, 15]. Allelic-dropout can be particularly problematic when genotyping tandem-repeat sequences because expanded repeats, known to cause disease, can be difficult to amplify. For example, the promoter of the fragile-X mental retardation gene, FMR1, contains a (CGG)_n_ repeat that can interfere with polymerase activity [16–18]. Expansion of this repeat causes fragile-X disease [19]. Recently, the Pacific Biosciences sequencer has demonstrated the ability to sequence these difficult repeats [18]

While these structures are of interest, both *in-vivo* and *in-vitro*, there is limited knowledge about which sequences have the potential to form non-B-DNA structures. Some of these structures have been studied in detail, and methods exist to predict the sequences in which they might form [20–24]. For example, the G-quadruplex structure has been well characterized, and methods exist to predict which sequences might form this structure [20–22]. However, these prediction methods are not definitive and some of these predicted structures may not be thermodynamically stable.

Uncertainty about which sequences have the ability to form non-B-DNA structures can make PCR primer design a challenge. Furthermore, if these structures have biological significance, predicting where they might exist could improve our understanding of genome function. If these structures are interfering with DNA and RNA polymerase *in-vivo*, then knowing where they may form in the genome will help us determine which regions could have reduced rates of transcription and/or DNA replication, such as those seen in [25, 26]. This may be of special importance for tandem repeat regions, where variation in tandem repeat length is common and can determine the stability of non-B-DNA structures [27]. Understanding how tandem repeats of various lengths can form non-B-DNA may help us better predict when, where and how they can cause disease.

## Detecting DNA structure with polymerase

Polymerase pausing is associated with non-B-DNA structures *in-vitro*, and polymerase stoppage assays are a common and relatively inexpensive method to detect potential non-B-DNA forming regions [12, 28]. Polymerase typically moves quickly along the DNA molecule, with the absolute speed depending on the system [13, 29]. When it encounters a non-B-DNA structure, such as a G-quadruplex or slipped strand structure, the polymerase pauses while the structure is resolved [12–14, 28, 30]. Polymerase stoppage assays can provide base-pair resolution for the positions at which the pauses occur, but only examine stoppage in bulk for a large number of DNA polymerase interactions [12, 28]. These assays do not provide realtime measurements of pausing, and do not measure the pausing at single-molecule resolution. Therefore, they can act as a way of measuring polymerase pausing on average, but do not detail the interaction between DNA and an individual polymerase.

Realtime detection of potential DNA structures can be accomplished with a nanopore device [31], or alternatively by using polymerase with a FRET-based approach [13]. However, these approaches have only been used for a small number of sequences, and the data are not readily available. Another method to detect DNA structure uses the realtime polymerase kinetics of the Pacific Biosciences sequencer [29, 32]. Here we outline a statistical approach by which polymerase kinetics can be measured and related to the nucleotide content of the template DNA at multiple scales using wavelets.

## Measuring polymerase kinetics

If polymerase pausing can indicate potential non-B-DNA structures, how should this pausing be measured? One method might measure the rate of polymerization along the sequence and search for regions of the sequence in which the polymerization rate was slow. Another method might examine the rate of change of polymerization, searching for regions in which the polymerase was slowing down. Both of these methods are encompassed with the wavelet approach developed here.

Wavelet transformations of a sequence produce two types of coefficients: smooth coefficients (*s*_*i*_) and detail coefficients (*d*_*i*_). The smooth coefficients are weighted sums, and the detail coefficients are the differences between these weighted sums [33]. A major benefit of the wavelet approach is that it produces these coefficients at different scales, with each scale increasing by a factor of two, allowing polymerase kinetics to be measured in regions of various sizes.

We examine multiple scales because we have limited knowledge about how polymerase interacts with non-B-DNA structures. The distance between the structure and the position at which the polymerase pauses is unknown. Furthermore, a structure might cause pausing at a specific location, but may also have large scale effects on polymerase kinetics by altering the surrounding sequence. For example, when viewed under an electron microscope, the regions around putative G-quadruplexes appear to form single-stranded loops [34]. Measuring the polymerase kinetics on multiple scales allows us to examine both fine-scale and course-scale interactions between the polymerase and the DNA template.

## Pacific Biosciences sequencer kinetics

The realtime polymerase kinetics of the Pacific Biosciences sequencer provide an unprecedented view of the interaction between polymerase and DNA. The sequencer follows a polymerase and its interaction with a single DNA molecule in a 100 nm well, detecting the timing of nucleotide incorporation with fluorescently labeled deoxyribonucleoside triphosphates [29]. This detailed recording of DNA synthesis allows modified nucleotides, such as 5-methylcytosine and N-6 methyladenine to be detected in realtime [35–40]. Potential non-B-DNA structures can also be detected realtime, but previously only hairpin structural regions have been examined [29, 32].

The analyses presented here examine the polymerase kinetics at each position in the genetic sequence. The kinetics are dominated by the interpulse duration (IPD, measured in seconds) between the adjacent nucleotide reads [29, 41, 42]. By measuring the IPD at each nucleotide, we examine how the kinetics are related to the sequence composition. Both sequence composition and kinetics are measured at multiple scales using the wavelet method.

A major benefit of using PacBio sequence data to uncover possible non-B-DNA regions is the volume of data produced in the sequencing process. The current system outputs a large amount of data per sequencing run, tens of thousands of reads with average read lengths of thousands of nucleotides. Sequencer kinetics have been made available online [35, 42], for example, from the sequencing of entire bacterial genomes for six species [35]. We use some of this available data in our analyses here, from the *E. coli* whole-genome amplification [42].

## Results and Discussion

### Correlations between nucleotide composition and kinetics at multiple scales

We begin our analysis by examining how the nucleotide composition is related to the kinetics. Using the wavelet analysis we examine these kinetics at multiple scales, for both the smoothed coefficients and the detail coefficients. We convert the read information into wavelet coefficients, for the nucleotide composition of the DNA template, IPD and for the number of incorrect nucleotides inserted (inserts) at each position in the aligned read. The inserts may represent actual insertion errors, but are more likely to represent sequencing errors [29]. The nucleotide smooth wavelet coefficients can be understood as the density of each nucleotide (A, T, C and G) at the varying scales, and the detail wavelet coefficients are the changes in this density.

Figure 1 shows the pairwise correlations between smooth wavelet coefficients (top right) and detail wavelet correlations (bottom left), with positive correlations highlighted in red and negative correlations highlighted in blue. The diagonal displays the relative power of the wavelet coefficients, indicating the strength of ability to test for correlations between the coefficients at that scale. For all of the coefficients examined here, the finest scales have the highest power.

**Figure 1.**
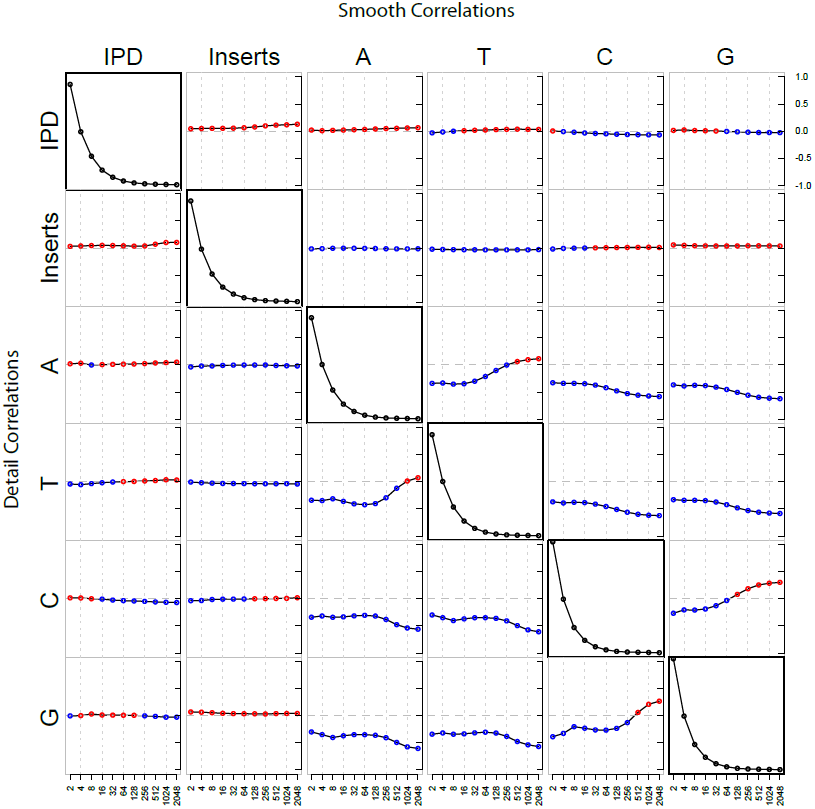
Correlations between wavelet coeffcients. The Pearson’s correlation between all of the wavelet coeffcients are shown: red indicating a positive correlation, blue a negative correlation. The correlation between detail coefficients is in the bottom left, and the correlation between smooth coefficients is in the top right. The power for the detail coefficients is in the diagonal. The frequency of each base in these reads is 25%. Only reads longer than 1,000 nucleotides were used to avoid complications caused by boundaries between reads. A total of 2^25^ nucleotides were examined (approximately 33.5 million nucleotides).

In Figure 1 the correlations between the kinetics and the nucleotide composition for the smooth coefficients are found in the top row, and for the detail coefficients in the leftmost column. These results indicate that only a small amount of the variation in kinetics can be attributed to the template nucleotide composition alone. Although some nucleotides are slightly faster or slower than others, the differences in their kinetics are not large.

These results also demonstrate the benefits of examining multiple scales; some of these correlations change as the scale changes. For example, the density of the nucleotide guanine is positively correlated with IPD at fine scales, but at scales above 32 nucleotides the correlation becomes negative (top right corner of Figure 1). In contrast, the change in the density of guanine is negatively correlated with the change in kinetics, but only at the finest and coarsest scales (bottom left corner of Figure 1).

However, these large-scale correlations must be interpreted carefully. The nucleotides are not evenly distributed throughout the genome and this can be seen in the correlations between the nucleotide wavelet coefficients. Because only one nucleotide can be attributed to each position in the genome, the correlations between each nucleotide are negative at the finest scales. As the scale increases we begin to see the nucleotide biases in the sequence: adenosine and thymine are positively correlated at large scales, and so are cytosine and guanine. This indicates that some regions of the reads are enriched with adenosine and thymine, and others enriched for cytosine and guanine. This approach is slightly different than measuring C/G (A/T) content, as the cytosines and guanines (adenosine and thymine) are enriched together on the same strand. These large-scale nucleotide biases could be partially responsible for the change in the correlations between kinetics and nucleotide composition between scales.

This method of sequencer read interpretation provides a novel way to view polymerase kinetics. By examining coefficients for regions of increasing size we find that some properties of the kinetics are scale variant. For example, the IPD and insert count are slightly positively correlated, and this correlation increases with the scale. Therefore, regions in which the polymerase moves slowly have slightly more insertion errors, and this becomes more pronounced as the region size increases. This method breaks the reads down into regions of varying size, but does not attribute values to individual nucleotides in the sequencer read. Therefore, it is only useful for examining correlations within the reads.

### Examining kinetics in a 128 nucleotide window

To examine how scaled values of the kinetics are related to individual nucleotides in the reference sequence, we use a slight modification of the standard wavelet approach (termed non-decimated wavelet transform, [43, 44]). This approach allows the wavelet coefficients to be attributed to specific locations in the read. To localize the scaled values, we have developed a method that weights each nucleotide with the values of the IPD of the previous nucleotides, including more of the previous nucleotides as the scale increases. We chose this approach because modified nucleotides primarily influence the polymerase as it approaches the nucleotide of interest [41, 42]. Similarly, as the polymerase approaches a non-B-DNA structure, we expect pausing to occur before the polymerase reaches the nucleotides that are involved in the structure.

We use this approach to examine specific patterns of interest and their surrounding region. These patterns are found at multiple places in the *E. coli* genome, and we aggregate these different regions together to produce a signature of the combined regions. We are not particularly interested in the regions surrounding these patterns, and would expect that combining their kinetics would result in a plot that appears similar to the average kinetics of the entire genome. In this way, we can visually compare the kinetics within the sequences of interest to their surrounding region.

Our primary interest is in the kinetics within the pattern of interest. While the immediate surrounding nucleotides are likely to influence these kinetics, at least slightly, we assume that if a non-B-DNA structure exists within these patterns, its existence does not depend on the surrounding regions. We examine these kinetics only at the finest scales: the raw IPD values, the 2 nucleotide smooth wavelet coefficients, and the 4 nucleotide smooth wavelet coefficients. The raw IPD values provide a picture of how the polymerase moves through these patterns at the single nucleotide scale. The smooth values provide an alternative way to examine if the region is slowing the polymerase kinetics, using measurements of consecutive IPD values within reads. Scaling is important when searching for pause regions. The polymerase may often pause at a specific nucleotide, but if these pauses coincide with faster polymerase activity in the surrounding region, then the regional speed of the polymerase might be normal. By combining the kinetics of consecutive nucleotides within a read, we can detect if a region slows polymerase activity.

**Figure 2.**
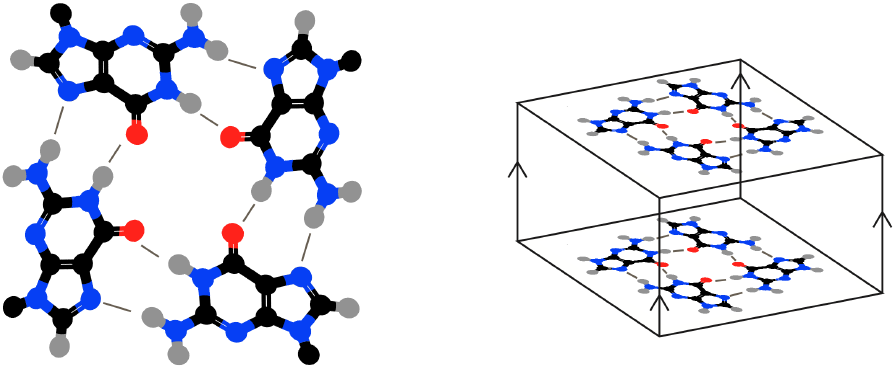
Guanine tetrad and parallel G-quadruplex. A tetrad formed by four guanines shown on left, with hydrogen bonds indicated by thin lines. These tetrads can stack to form G-quadruplexes, shown on the right. The G-quadrupluex shown has all of its strands in parallel, as indicated by the arrows connecting the two tetrads. (Image not to scale.)

### Kinetic patterns around (GGC)_n_ repeats

Tandem repeats composed of (GGC)_n_ have the potential to form non-B-DNA structures, but the structures that they can form depend on their length ([26, 45–47], reviewed in [48]). Short (GGC)_n_ repeats can form stable quadruplexes (e.g., Figure 2), while longer repeats form a hairpin-like structure with some similarity to the G-quadruplex structure [26, 45–47]. These structures may form *in-vivo*, impeding DNA replication [26, 49]. Expanded (GGC)_n_ repeats are difficult to genotype because of their impact on polymerase activity [16, 17, 30, 49–51] but these repeats can be sequenced with Pacific Biosciences technology [18]. Intriguingly, the Pacific Biosciences sequencer kinetics of the fragile-X repeats have strand asymmetry. When the template is (GGC)_n_, the polymerase is slower, and has a higher variance, than when the template is (CCG)_n_ [18], as would be expected if the G-rich template is forming a non-B-DNA structure.

Here we examine the kinetics of the sequence (GGC)_3_GG, which can be found at 27 locations in the *E. coli* genome. The *E. coli* genome does not contain long (GGC)_n_ repeats, although the repeats examined here have the ability to expand, and some strains of *E. coli* contain expanded (GGC)_n_ repeats [52]. This sequence is capable of forming a G-quadruplex structure with two stacked guanine tetrads (Figure 2). These G-quadruplexes can take various forms, but require the planar guanine tetrads to coordinate. If the strands of the quadruplex are found in parallel (as seen in Figure 2) then the guanines are in the *anti* position, just as they are in normal B-DNA. However, if any of these strands run anti-parallel, some of the guanines must be in the *syn* position, up-side down from their normal orientation [2]. The effect that *syn* nucleotides have on polymerase kinetics is unknown.

**Figure 3.**
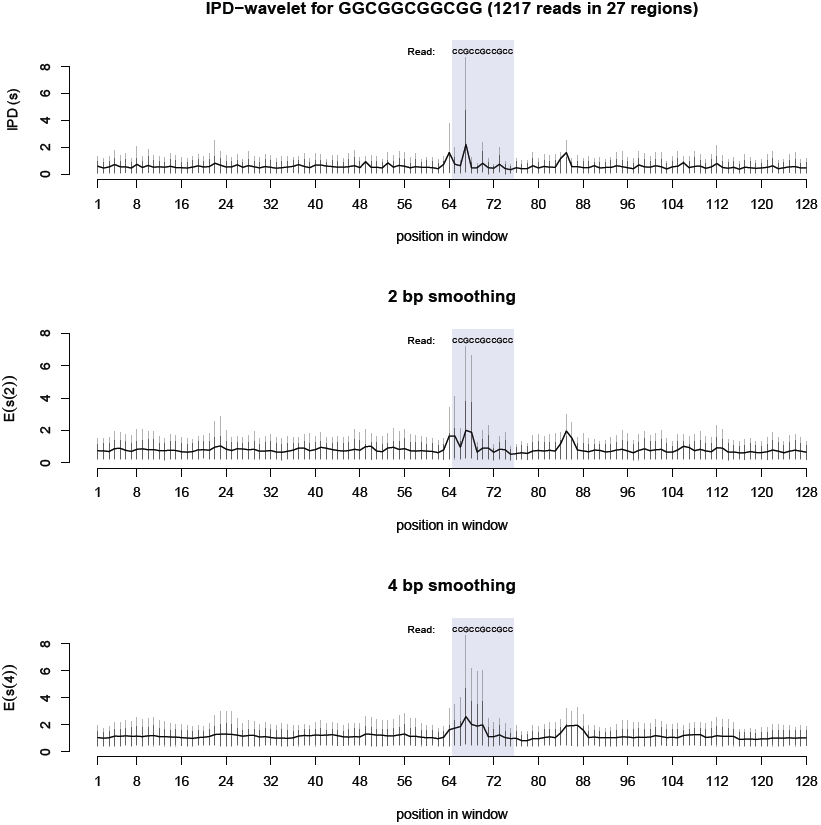
Kinetics around CGG repeats. These three graphs show the average polymerase kinetics for the sequence GGCGGCGGCGG, found in 27 regions in the *E. coli* genome. A total of 1217 reads were used in the analysis. The top graph shows the raw IPD values for the region, in seconds. In the middle and bottom graphs, the 2 bp (4 bp) smoothing represents wavelet smoothing over a two (four) nucleotide region within the reads. For each measurement, the black vertical lines are the 90% quantile, the grey vertical lines are the 95% quantile.

The kinetics around this repeat sequence suggest that a non-B-DNA structure is present (Figure 3). Pauses occur just before the sequence, and then again three times within the sequence. All of the pauses occur at the base immediately preceding the pairs of guanine. If a G-quadruplex were present in these reads, the cytosines would form the “loops” of the G4 structure. These pauses are striking because of their magnitude. At the slowest position, the first cytosine in the sequence, 10% of the reads have an IPD value of four seconds or greater, and 5% have an IPD greater than 8 seconds. In comparison, the IPD in most of surrounding region rarely has IPD values over 1 second.

Using wavelet smoothing, the cumulative speed of the polymerase in the region can be measured. The smoothed wavelet coefficients represent the weighted sums within each read for the nucleotide and the nucleotides preceding it. The smoothed results here indicate that the region slows the polymerase. For most of the surrounding sequence, the smoothing removes the variation in the kinetics. However, there is a pause region around nucleotide 85 in the window that may represent another structure present in one or more of the regions examined. The regions surrounding these repeats were not examined further.

Importantly, the pausing here is strand specific. The opposite strand, composed of (CGG)_3_CC, does not present any impedance to the polymerase and the kinetics in this sequence are nearly indistinguishable from the surrounding region (Figure 4). This rules out C/G content or presence of CpG dinucleotides as the sole cause of the pausing around the (GGC)_n_ repeat region.

Without further experimentation, we cannot conclude that G-quadruplexes are present in these reads with certainty. However, the pattern of kinetics around the sequence examined appear strikingly similar to what would be expected if a G-quadruplex were occurring. Further work will be necessary to determine whether these pauses are in fact quadruplexes, or whether another structure is present. Nevertheless, something is causing the polymerase to slow down in the region, suggesting a non-B-DNA structure is present.

**Figure 4.**
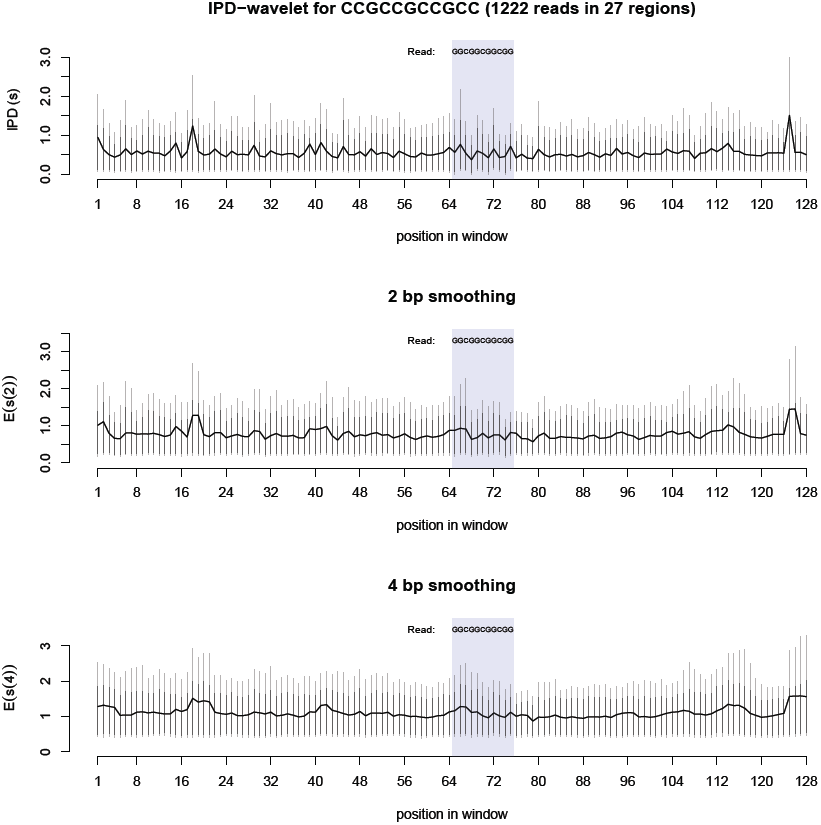
Kinetics around CCG repeats. These three graphs show the average polymerase kinetics for the sequence CCGCCGCCGCC, found in 27 regions in the *E. coli* genome. A total of 1222 reads were used in the analysis. The top graph shows the raw IPD values for the region, in seconds. In the middle and bottom graphs, the 2 bp (4 bp) smoothing represents wavelet smoothing over a two (four) nucleotide region within the reads. For each measurement, the black vertical lines are the 90% quantile, the grey vertical lines are the 95% quantile.

### Kinetic patterns around (CG)_n_ repeats

Z-DNA is a left-handed double helix that, roughly speaking, can form in alternating purine-pyrimidine sequences with a G/C content above 50% ([23, 24] reviewed in [53]). Z-DNA has a zig-zagging backbone, and its guanines are found in the *syn* position, forming Hoogsteen base-pairing with their corresponding cytosines [53]. Z-DNA can be induced in these sequences if they are under torsional strain [3], even in minute amounts [54]. Z-DNA can cause transcriptional blockage, but only under torsional strain [55].

The tandem repeat (CG)_n_ is the most stable Z-DNA forming sequence, and longer repeats are more likely to form Z-DNA [53]. However, repeats as small as (CG)_3_ have been shown to form Z-DNA [56]. The *E. coli* genome contains 9 regions with the repeat (CG)_5_. Because the repeat is palindromic, there are 18 different unique regions with this pattern (9 forward strand, 9 reverse strand). As before, we aggregate these regions together to examine their kinetics.

The kinetics within this repeat produce an interesting pattern (Figure 5). Most of the cytosines in the repeat region have a slightly high average IPD value, while the guanines have a slightly lower average IPD value. More interesting, the cytosines can sometimes have an IPD over 1 second. As indicated by the vertical bars in Figure 5, 10% of the IPD values at most of the cytosines are over 1 second, and 5% are greater than approximately 2 seconds in duration. In contrast, the majority of the IPD values at the guanines are very small (most of their 90% and 95% ranges are less than 0.5 seconds).

Though interesting, these results are not strong evidence that any non-B-DNA structures occur in these repeat regions. This region does not cause obvious polymerase pausing, as indicated by the smoothed wavelet values. The alternating high and low IPD values suggests that the backbone may be in a unique formation and/or some of the guanines may be in *syn*. However, most sequences have a unique kinetic signature [32, 41, 42] and these results might simply be the signature of this sequence in its B-DNA form.

Z-DNA might partially account for the kinetics seen here. However, if Z-DNA is having an effect, it is more likely to have been present before the polymerase reached the repeat region, leaving some or all of the guanines in the *syn* position and the DNA backbone in a zig-zagging form. Therefore, these kinetics might not represent Z-DNA, but rather an intermediate structure that has a unique kinetic signature. Further experimentation might be able to stabilize Z-DNA within the sequencer, helping to determine if and how it affects polymerase kinetics.

### Potential applications

Knowing which sequences can pause polymerase can be useful, especially for tandem repeats. In PCR, allelic dropout can cause genotyping errors if a heterozygote has one allele that causes pausing and another that does not. Alleles that replicate slowly can be out-competed by faster alleles. If a guanine rich tandem repeat is the target, this allelic dropout can be especially problematic [15]. By predicting which sequences cause polymerase pausing, methods can be employed to avoid allelic-dropout in PCR amplification [15, 57, 58].

**Figure 5.**
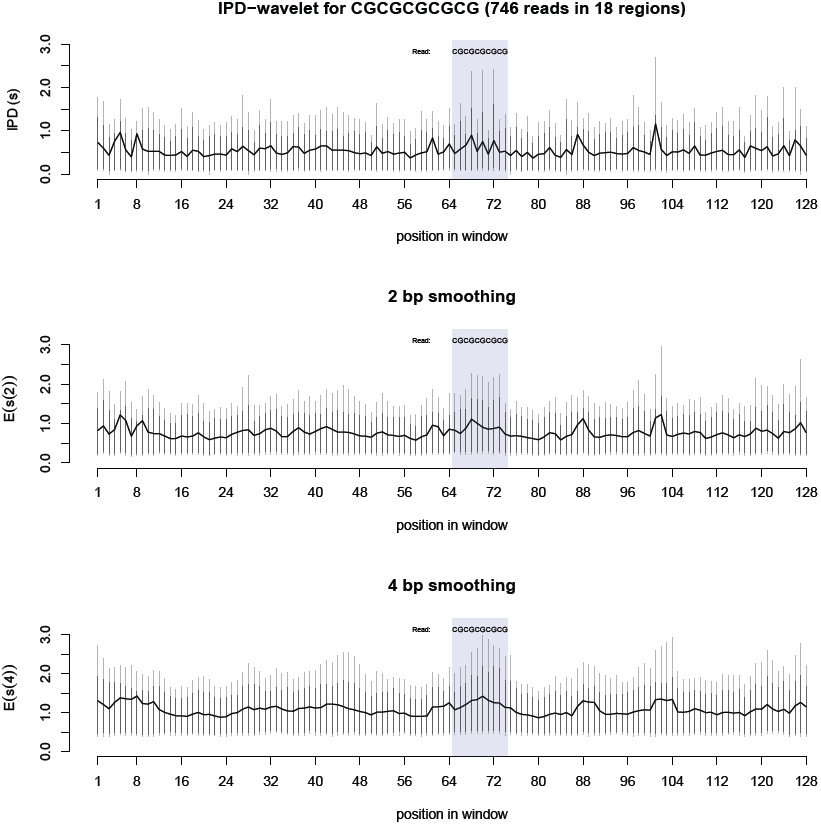
Kinetics around CG repeats. These three graphs show the average polymerase kinetics for the sequence CGCGCGCGCG, found in 18 regions in the *E. coli* genome. A total of 746 reads were used in the analysis. The top graph shows the raw IPD values for the region, in seconds. In the middle and bottom graphs, the 2 bp (4 bp) smoothing represents wavelet smoothing over a two (four) nucleotide region within the reads. For each measurement, the black vertical lines are the 90% quantile, the grey vertical lines are the 95% quantile.

Even if PCR is not used, polymerase pausing may nevertheless introduce genotyping errors. If one allele causes a large amount of pausing in the Pacific Biosciences sequencer, then its coverage will be reduced. If coverage is low, then an allele that pauses might appear as a sequencing error because it will be covered at a lower frequency than alleles that do not cause pausing. The extent to which this will be a problem for sequencing has yet to be determined, but relative rates of sequencing should be considered as a potential source of sequencing errors, especially in tandem repeat sequences where changes in tandem repeat number can alter non-B-DNA forming potential [27].

With the large amount of Pacific Biosciences sequencer kinetics available, new pause sites can be uncovered. These pause sites might be regions of non-B-DNA formation, and could be targeted for further research on non-B-DNA structures. While the exact structure that forms in the sequencer may never be determined, the important fact remains that these sequences can cause DNA polymerase pausing, and this has implications for PCR as well as our understanding of genome function. If RNA polymerase pauses at these regions in a similar manner, then the pausing seen in the sequencer might indicate regions where DNA has an intrinsic ability to reduce gene expression, as has been found for G-quadruplex sequences [59, 60]. If the DNA structure is acting as a polymerase speed-regulator [25], then some DNA sequences would be intrinsically more difficult to transcribe and/or replicate. This has the greatest implication for regions with tandem repeats, because tandem repeat variation is common due to the high rate of tandem repeat expansion and contraction, and these common polymorphisms can result in variation in DNA structural potential [27].

Human promoters are enriched with sequences predicted to have G-quadruplex forming potential [61]. Promoter G-quadruplexes are associated with transcriptional pausing [60, 62] and appear to affect relative rates of gene expression [63, 64]. Some of these promoter G-quadruplex regions are composed of tandem repeats [65]. When these quadruplexes are on the template strand, they have the ability to pause RNA polymerase and reduce the rate of transcription. If a tandem repeat with quadruplex forming potential were to expand, it could potentially result in an allele that is nearly incapable of being transcribed. By examining how tandem repeats cause polymerase pausing in Pacific Biosciences sequencer reads, we might better predict which tandem repeats impede gene expression.

Ultimately, the ability of a sequence to pause polymerase may be directly related to mutation rates. Secondary structures are related to DNA mutation, especially around tandem repeats [52, 66]. In *E. coli*, the formation of DNA hairpins is associated with tandem repeat instability on the leading strand of DNA synthesis [52] and non-B-DNA forming sequences can increase the local rate of mutation [66]. By improving our ability to predict which sequences form non-B-DNA structures, we may better predict where mutations will occur.

And finally, research into how DNA structures can pause polymerase in the Pacific Biosciences sequencer may be useful for predicting modified nucleotides. If different structures are present in different reads, then accounting for these different states could improve detection of modified nucleotides. This is especially interesting considering that nucleotide modification can influence non-B-DNA structural formation and stability. For example, Z-DNA is stabilized by cytosine methylation [67–69]. Modified nucleotides were not present in the DNA that was used in this analysis.

The wavelet model used here has the potential to be developed further to allow computationally efficient modeling of the kinetics at multiple scales. Regions which show polymerase pausing can then be targeted for more intricate models of the potential DNA conformations. Our analyses here suggest that these polymerase kinetics provide an unprecedented view of DNA structure, and as more Pacific Biosciences data becomes available, methods examining non-B-DNA structure can be refined. Furthermore, these methods can also be applied to other approaches, such as the FRET based approach to measuring polymerase speed [13] or nanopore based detection of non-B-DNA structure [31]. We hope these and future results will improve our understanding of DNA chemistry and its relationship to polymerase activity. To aid other researchers studying Pacific Biosciences kinetics, we have released a package for R on GitHub that allows the analyses presented here to be repeated for any DNA pattern of interest on any Pacific Biosciences kinetics data (see methods).

## Methods

### Data

Sequencing data from the whole genome amplification of *E. coli* genome were used, taken from the supporting information of [42]. These data are currently available from Pacific Biosciences at: http://www.smrtcommunity.com/Share/Datasets Native DNA (untreated DNA from the *E. coli* genome) was not used in our analyses here because modified nucleotides present in native DNA affect kinetics and have an unknown effect on alternative DNA structure formation.

### Decimated wavelet transform

We examine properties of a sequence, *S*, with a length *N* (i.e., *S* = {*s*_1_, *s*_2_, *s*_N_}). To simplify the wavelet approach we assume that N is a power of two, but modified wavelet approaches permit sequences of any length [33]. A decimated wavelet decomposition of the sequence results in a set of coefficients for each scale, *j*, where *j* ∊ {1, 2,…, *log*_2_(*N*)} = *J*. These coefficients take two forms, the smooth coefficients, 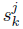, and detail coefficients, 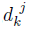. The smooth coefficients are generated with a low pass filter, 𝓗, and the detail coefficients are generated with a high pass filter, 𝒢, applied recursively to generate coefficients for each scale [33, 70]. The equations for the low and high pass filters can be written as:

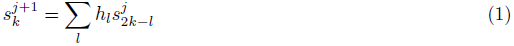

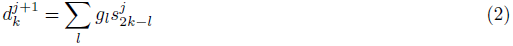

For *k* = 1,2,…, *N*/2^*j*^, where *h* and *g* represent the wavelet filter used to transform the sequence. For this study we chose the Haar wavelet filter [71]. The Haar wavelet is the simple and easily understood. Additionally, the Haar wavelet can easily detect sharp changes that occur in the sequence [33]. For the Haar wavelet, 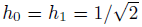 and 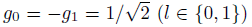. For clarity, we will denote each scale of each coefficient by the size of its region, so our scales are {(2), (4),(*N*/2)} (e.g., coefficients at the 2 base-pair scale are denoted as 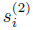 and 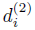). For an example, we apply the low pass filter (1) and high pass filter (2) to produce the smooth and detail coefficients for the 2 nucleotide scale using the first positions:

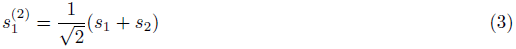

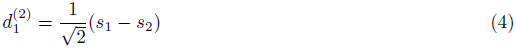

To find the coefficients at higher scales, the filters are applied to the coefficients of the previous scale. For example:

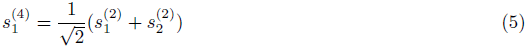

The number of coefficients at each scale is half of that of the previous scale. Therefore, the decimated wavelet coefficients can be attributed to regions of size *j*, but not individual nucleotides in the sequence. This has the disadvantage of being shift-variant, so that if the sequence is shifted by one nucleotide, the wavelet coefficients will change. However, we do not believe that shift-variance is an issue for our analysis here. We generate coefficients within the reads for: nucleotide composition, polymerase kinetics and insert error. The reads do not all begin at the same place in the sequence, so the wavelet coefficients do not all originate from the same place in the sequence. Therefore, this approach that is nearly shift-invariant in the DNA sequence, but not in the read.

A major benefit of decimated wavelets is the orthonormality of the wavelet basis, assuming the appropriate wavelet coefficients are used [70, 72]. This allows us to examine variation specific to each scale and also to independently examine relationships between different sequence properties at different scales. This approach is used in signal processing for compression of video and audio signals, and is often referred to as multi-resolution analysis [33]. Wavelets have also been used to analyze properties of DNA and protein sequences [65, 73–76], for example examining how different elements in the genome are related [65, 73].

We limit our analysis to reads longer than 1000 nucleotides and concatenate the reads together, trimming the resulting vector to its nearest power of two (here 2^23^ nucleotides). Some of the resulting wavelet coefficients span the boundaries between reads, but we do not feel the low number of these discontinuities create a problem for the resulting analysis.

All analyses were performed using custom R scripts [77], using the package Wavethresh [78].

### Wavelet interpretation of DNA nucleotides

We convert DNA sequences into a wavelet interpretation so that we can compare wavelet coefficients of the sequence kinetics with nucleotide composition of the read. Nucleotide composition can be seen as a vector in which the nucleotides of interest have a value 1 and all other positions have a value of 0. For example, the sequence 5’-ACTG-3’ can be seen as a set of four vectors for each nucleotide: *A* = {1,0,0,0}, *C* = {0,1,0,0}, *T* = {0,0,1,0}, *G* = {0,0,0,1}). Each of these vectors can then be transformed with the wavelet decomposition to form wavelet coefficients for each nucleotide. The same approach can be used to examine any sequences of interest. We examine correlations between the wavelet coefficients for each factor using Pearson’s product moment correlation.

The power of the wavelet coefficients at any scale is measured as the relative contribution of the sum of the squares of the detail coefficients at that scale to the total sum of squares:

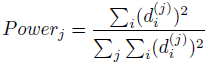

### Non-decimated wavelet transform

The shift-variance of the decimated wavelet transform becomes an issue when we want to attribute wavelet coefficients to specific nucleotides in the sequence. To overcome this issue, a non-decimated wavelet transform was used for the 128 nucleotide window analysis [43, 44]. The non-decimated wavelet produces wavelet coefficients at each position in the sequence by using a decimated wavelet transform on each possible shift. The result is a set of wavelet coefficients representing scaled values at each nucleotide for each scale. The resulting wavelet basis is no longer orthonormal, although orthonormal sets can be extracted [43, 44]. The resulting values are similar to those computed with the decimated approach above, because this approach is decimated in the sequencer reads not the DNA sequence. The difference here being that coefficients are generated for all possible shifts in the sequence, and then attributed to specific nucleotides in the reference sequence.

We modify the standard approach slightly here so that the results are more applicable to polymerase kinetics. The non-decimated approach attributes each wavelet coefficient to the position in which that coefficient starts. This results in a value that is, for the Haar basis, the weighted sum of the value at that position and the values at the downstream nucleotides. If we used this method to measure polymerase kinetics, the scaled rate of polymerase activity at a specific nucleotide would depend on the rate at nucleotides that have yet to be sequenced. So for all positions *i* in the sequence, the non-decimated high pass filter for the 2 nucleotide scale would be:

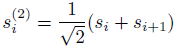

This would attribute pauses to positions upstream of the pause site. This is the opposite of the current model of Pacific Biosciences polymerase kinetics, in which modified nucleotides primarily influence the kinetics before the modified nucleotide is sequenced, as it enters the polymerase, [41, 42]. If we are to stipulate that a DNA structure might influence the kinetics as the polymerase approaches the structure, and not after the structure has been sequenced, then we should attribute pauses to the nucleotides at which they occur and the region *downstream* of these nucleotides. This is achieved here by simply reversing the sequence before it is transformed. Following from the example above, if we reverse the sequence before transformation, the coefficient becomes:

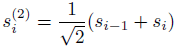

The lack of symmetry of the transform is beneficial in this case, but is not entirely necessary for all wavelet approaches. The Daubechies least-asymmetric transforms can be used in cases for which greater symmetry is desired [70].

### R package for analyzing Pacific Biosciences data

The analyses presented here can be reproduced for any Pacific Biosciences data using an R package that can be found with installation instructions at: https://github.com/sterlo/kineticWavelets This package provides a general method for analyzing Pacific Biosciences polymerase kinetics for any DNA pattern of interest, and allows DNA patterns to be defined using regular expressions. The use of regular expressions provides flexibility in a DNA pattern definition. This flexibility is useful for examining sequences predicted to form various DNA structures, because the patterns predicted to form various structures can be highly variable. For example, a stable G-quadruplex is often defined as: (*GGG*)(*N*)_1-7_ (*GGG*)(*N*)_1-7_(*GGG*)(*N*)_1-7_(*GGG*) which can be defined with the regular expression: “(GGG(.){1,7}GGG(.){1,7}GGG(.){1,7}GGG)”

We hope this package will be used to analyze other potential DNA structures in Pacific Biosciences data with the wavelet approach used here.

## Acknowledgements

The authors are grateful to Steve Turner and Jonas Korlach, founders of Pacific Biosciences Inc. for their helpful discussions and insights into the polymerase kinetics of their sequencer.

